# Investigating Transfusion-Related Sepsis using Culture-Independent Metagenomic Sequencing

**DOI:** 10.1101/653337

**Authors:** Emily Crawford, Jack Kamm, Steve Miller, Lucy M. Li, Saharai Caldera, Amy Lyden, Deborah Yokoe, Amy Nichols, Nam K. Tran, Sarah E. Barnard, Peter M. Conner, Ashok Nambiar, Matt S. Zinter, Morvarid Moayeri, Paula Hayakawa Serpa, Brian C. Prince, Jenai Quan, Rene Sit, Michelle Tan, Maira Phelps, Joseph L. DeRisi, Cristina M. Tato, Charles Langelier

## Abstract

**Background:** Transfusion-related sepsis remains an important hospital infection control challenge. Investigating septic transfusion events is often restricted by the limitations of bacterial culture in terms of time requirements and low yield in the setting of prior antibiotic administration.

**Methods:** In three Gram-negative septic transfusion cases, we performed mNGS of direct clinical blood specimens in addition to standard culture-based approaches utilized for infection control investigations. Pathogen detection leveraged IDSeq, a new open-access microbial bioinformatics portal. Phylogenetic analysis was performed to assess microbial genetic relatedness and understand transmission events.

**Results:** mNGS of direct clinical blood specimens afforded precision detection of pathogens responsible for each case of transfusion-related sepsis, and enabled discovery of a novel *Acinetobacter* species in a platelet product that had become contaminated despite photochemical pathogen reduction. In each case, longitudinal assessment of pathogen burden elucidated the temporal sequence of events associated with each transfusion-transmitted infection. We found that informative data could be obtained from culture-independent mNGS of residual platelet products and leftover blood specimens that were either unsuitable or unavailable for culture, or that failed to grow due to prior antibiotic administration. We additionally developed methods to enhance accuracy for detecting transfusion-associated pathogens sharing taxonomic similarity to contaminants commonly found in mNGS library preparations.

**Conclusions:** Culture-independent mNGS of blood products afforded rapid and precise assessment of pathogen identity, abundance and genetic relatedness. Together, these challenging cases demonstrated the potential for metagenomics to advance existing methods for investigating transfusion-transmitted infections.

## Introduction

While transfusion-associated infections have globally decreased over the past 50 years, bacterial contamination of the platelet supply remains a significant public health challenge, with approximately one in 1,500 – 5,000 units containing detectable organisms[1–5]. Room temperature storage facilitates more bacterial growth in platelets compared to other blood products, with skin flora, asymptomatic donor bacteremia, and direct introduction from environmental sources accounting for the majority of contaminants[3,6]. Even though current culture-based screening approaches may miss a significant fraction of contaminated units, clinically reported sepsis occurs in only one in 15,000-100,000 transfusions, presumably due to frequent concurrent antibiotic administration, low bacterial inoculum, or attribution of transfusion-related outcomes to a patient’s pre-existing infection or illness[3,7].

Pathogen-reduction of platelet products using amotosalen, a nucleic acid cross-linking agent activated by ultraviolet (UV)-A irradiation, has been routinely performed in Europe for over a decade and recently received United States Food and Drug Administration approval[8]. Photoinactivation is broadly effective against diverse bacterial pathogens, and while failure at high bacterial loads has been described, few reports of sepsis have been reported following treatment[9,10].

Here we describe three transfusion-related Gram-negative sepsis investigations including one involving a pathogen-reduced platelet product. For each, we performed metagenomic next generation sequencing (mNGS) in addition to standard microbiologic diagnostics and found that mNGS enabled rapid and precise taxonomic identification and phylogenetic analysis of bacterial isolates, as well as culture-independent assessment of direct clinical samples.

## Methods

### Ethics Statement

Investigations were carried out according to a no-subject-contact study protocol approved by the UCSF Institutional Review Board (IRB# 17-24056) which permitted analysis of de-identified leftover clinical microbiology samples from collaborating institutions and subsequent review of study subjects’ electronic medical records. No decisions regarding antibiotics or other patient-specific treatment interventions were made using sequencing data.

### Microbial culture

Patient blood cultures were performed via inoculation into BD Bactec Plus Aerobic and Lytic Anaerobic media (Becton Dickinson). This same approach was used for residual platelet transfusion cultures in cases one and three, and for residual red blood cell segment cultures in case 2. Species identification was performed using MALDI-TOF mass spectrometry (Bruker). Details on culture time to positivity are described in **Supplemental Table 1**. Sterility testing was carried out as described in the **Supplemental Methods**.

### Sequencing

Sample processing and DNA extraction were carried out as described in the **Supplemental Methods**. Ten to 100 ng of DNA from each sample was sheared with fragmentase (New England Biolabs) and used to construct sequencing libraries with the NEBNext Ultra II Library Prep Kit (New England Biolabs). Adaptor ligated samples underwent amplification with dual unique indexing primers. Libraries were quantified, pooled, and underwent paired end 150 base pair sequencing on an Illumina MiSeq or NextSeq 550. **Supplemental Table 2** lists the number of reads obtained for each sample.

### Bioinformatics and phylogenetic analyses

Detection, taxonomic identification, and abundance quantitation of microbes from raw sequencing reads was first performed using the IDseq pipeline according to described protocols[11]. To control for background environmental and reagent contaminants, no-template water control samples were incorporated alongside extracted nucleic acid and carried forward throughout library preparation and sequencing. Genome assembly was performed by first trimming the raw sequencing reads in fastq files using TrimGalore[12] and assembling using Unicycler[13] with default parameters. To identify the closest related species in cases one and two, BLAST+[14] against the National Center for Biotechnology Information (NCBI) nt database and the Mash/Mini Hash search via PATRIC[15,16] were employed to analyze assembled and annotated contiguous sequences (contigs). For case three, sequence type was determined using SRST2[17]; other *Klebsiella pneumoniae* sequences in NCBI databases belonging to the same sequence type were found using a Mash/Mini Hash search via PATRIC[15,16].

Phylogenetic analysis for case one and case three was used to determine the relationship between the samples in this study and the most closely related NCBI genomes. First, trimmed reads were aligned against a reference genome (case one: *A. baumannii* CP017642.1, case three: *K. pneumoniae* CP015392.1) using Snippy v4.3.6[18]. Then a SNP alignment was obtained by variant calling using bcftools v1.9[19]. We filtered SNPs with QUAL<30, and filtered genotypes with major allele depth (FMT/AD) < 10 or major allele frequency (FMT/AF) < .9. Finally, the maximum likelihood phylogeny based on the SNP alignments was built in RAxML v8.2.12[20] using ‘-m ASC_GTRCAT --asc-corr=lewis’ options.

## Results

### Investigation #1: *Acinetobacter* septic platelet transfusion

Patient A, a 59 year old man with relapsed acute lymphoblastic leukemia undergoing chemotherapy, received two doses of pathogen-reduced apheresis platelets on the day of planned home discharge. No side effects were experienced after transfusion of the first platelet unit, however after receiving the second unit two hours later, he developed chills and rigors followed by fever and critical hypotension. He received fluids and vancomycin before being transferred to the intensive care unit where meropenem, vasopressors, respiratory support and continuous renal replacement therapy were administered. He recovered fully and was discharged home 16 days later on parenteral antibiotics.

Blood cultures obtained 4.5 hours post-transfusion returned positive for *Acinetobacter calcoaceticus/baumanii (ACB) complex*. Residual material obtained from a saline rinse of the returned platelet bag also grew *ACB complex* as well as *Staphylococcus saprophyticus*. Culture of a non-transfused platelet co-component from the same donor, concurrently pathogen-inactivated with the transfused product five days earlier, was negative. At the time of the investigation, hypotheses considered included failure of the pathogen-reduction process, contamination of the pathogen-reduced platelet bag during transport, or pre-existing sub-clinical bacteremia in the patient.

To investigate this case further, we first sequenced the cultured *ACB* complex isolate from the patient’s blood and the *ACB* complex isolate from a wash of the transfused platelet bag. In total, sequencing and analysis for this case was performed in 45 hours. *De novo* genome assembly revealed a novel *Acinetobacter* species most closely related (92% average nucleotide identity) to *A. nosocomialis* and *A. pitii*, members of the *ACB* complex (**Figure 1A**). Phylogenetic analysis revealed zero single nucleotide polymorphisms (SNPs) over the 3.9 Mb genome between the novel *Acinetobacter* species cultured from the blood and from the platelet bag, indicating that the two isolates were identical.

**Figure 1.**
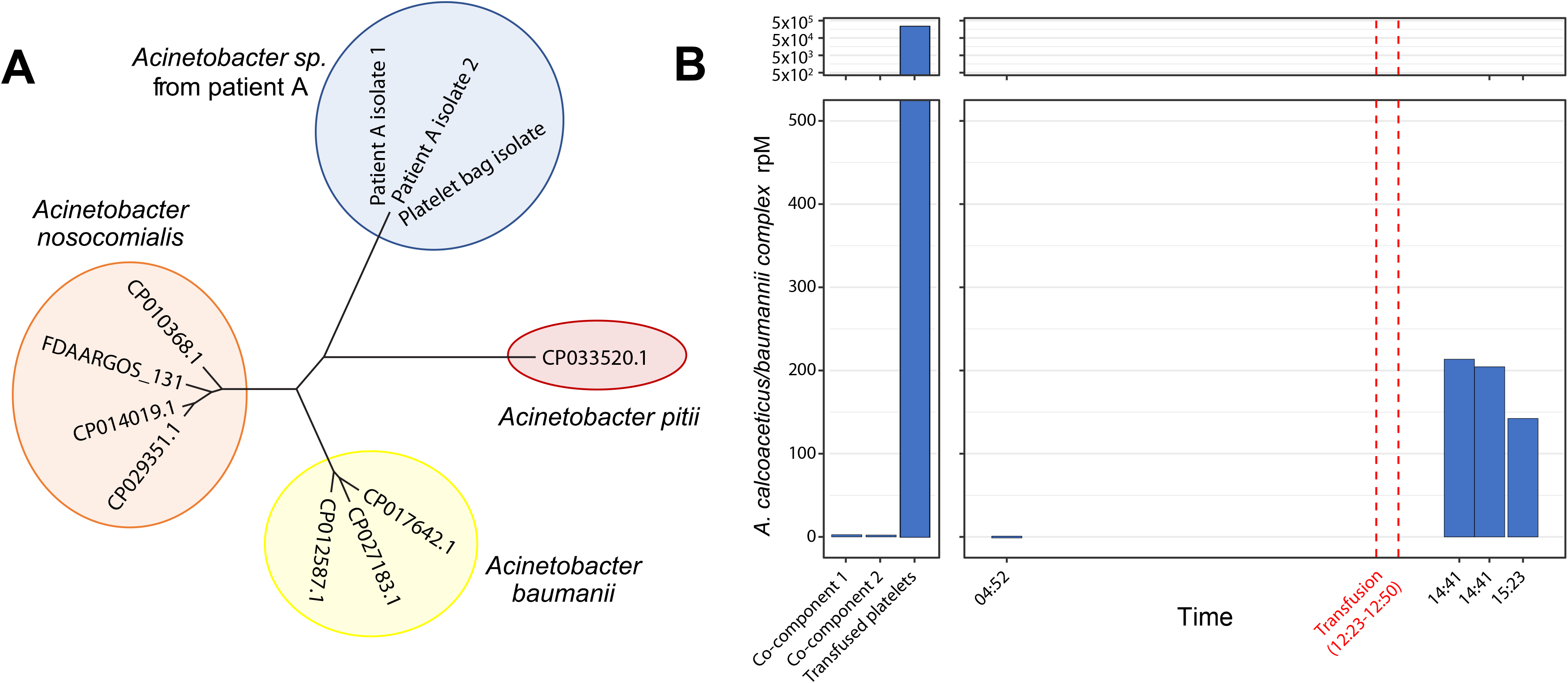
*Acinetobacter* septic transfusion investigation. **A)** Maximum likelihood phylogenetic tree based on SNP alignments demonstrates relatedness of the novel *Acinetobacter species* isolated from both patient A and the residual transfused platelet product relative to closely related species within the *Acinetobacter calcoaceticus/baumannii* (ACB) complex. **B)** Abundance of *ACB* complex in the transfused platelet product and co-components (left panel) and in patient A’s plasma (right panel), determined by culture-independent metagenomic sequencing and measured in reads per million, rpM.

Culture-independent mNGS detected a high abundance of *ACB complex* in the transfused product (230,000 reads per million, rpM) and in the patient’s plasma following transfusion (190 rpM) (**Figure 1B**), as determined by summing the reads found by IDSeq aligning to species within the ACB complex (NCBI taxid 909768) which includes *A. calcoaceticus, A. baumanii, A. nosocomialis* and *A. pitii* among others. By comparison, the untransfused co-components and patient’s plasma before transfusion had very few alignments to *ACB complex*, in the range of expected background levels (0.25 and 2.2 rpM, respectively). Assessment of these low abundance alignments at the species level revealed that they most likely represented mis-assigned reads from other *Acinetobacter* species, notably *A. johnsonii*. More specifically, the co-components and pre-transfusion samples had only a small percentage of *Acinetobacter* genus reads that best aligned to an *ACB species*, while in the transfused product and post-transfusion samples, the vast majority of genus *Acinetobacter* reads aligned to *ACB species* (**Supplemental Figure 1A**). The orders-of-magnitude difference in percent reads mapping to *ACB complex*, along with the change in composition of *Acinetobacter* assignments between samples, indicated that only the transfused product and patient’s post-transfusion plasma were contaminated by a species within the *ACB complex*. Although the sequence of the cultured isolate in this case allowed us to properly identify the strain phylogenetically, it was not required to complete the mNGS analysis that distinguished pathogen and environmental contaminant.

### Investigation #2: Fatal *Pseudomonas aeruginosa* septic transfusion

Patient B, a 77 year old man, was admitted for management of acute on chronic heart failure and implantation of a right ventricular assist device. During surgery, he received three units of platelets, three units of fresh frozen plasma, and two units of cross-matched packed red blood cells (PRBCs). Intraoperatively, the patient became hemodynamically unstable. Cultures drawn three hours post-surgery ultimately returned positive for *Pseudomonas aeruginosa*, raising concern for a septic transfusion reaction (**Figure 2**).

**Figure 2.**
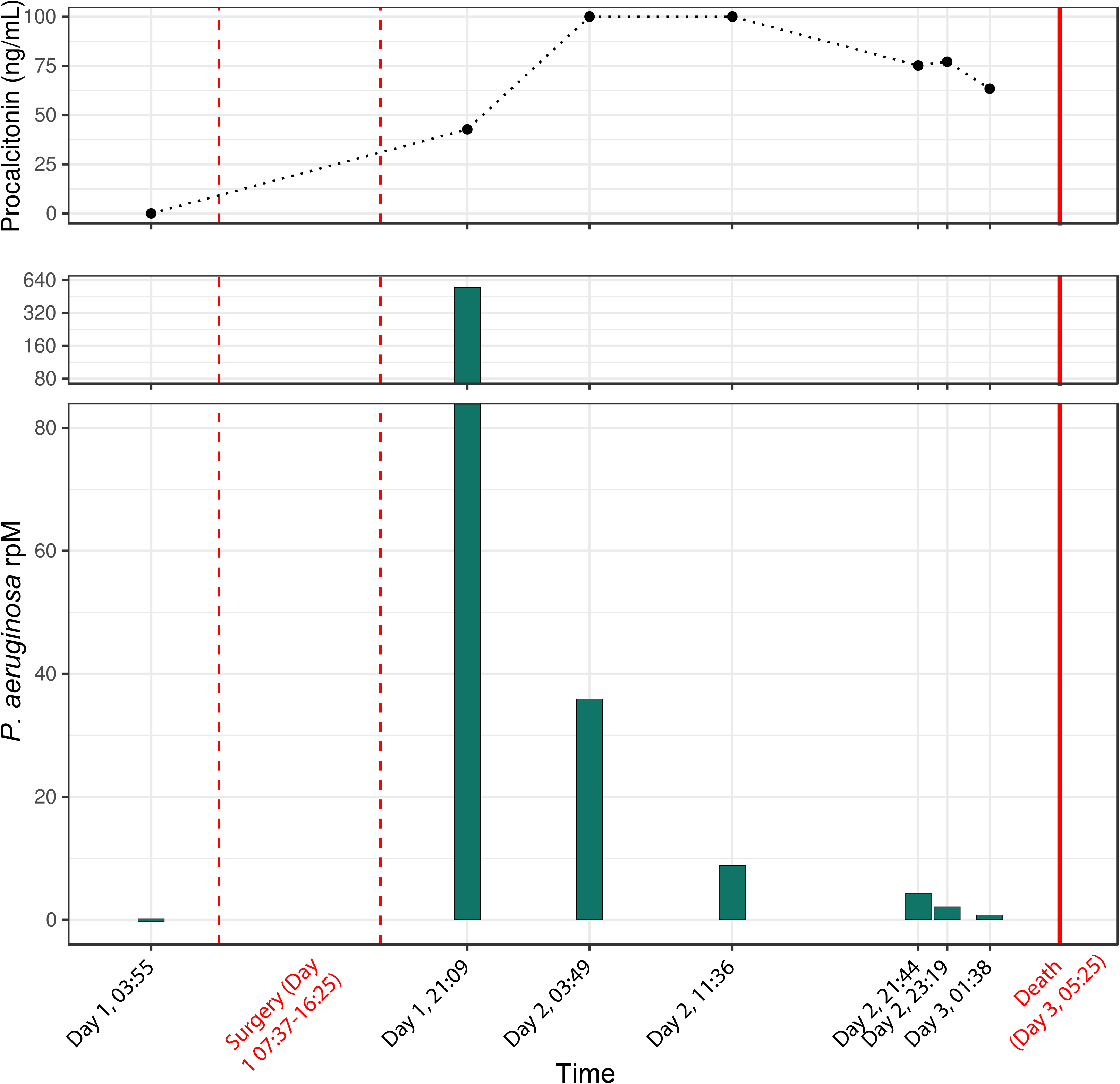
*Pseudomonas* septic transfusion investigation. Abundance of *Pseudomonas aeruginosa* in patient B’s plasma throughout the course of the fatal septic transfusion event, determined by culture-independent metagenomic sequencing and measured in reads per million (rpM, lower panel). Procalcitonin level (ng/mL) over the course of the septic transfusion event is plotted in the upper panel.

Despite administration of antipseudomonal antibiotics, the patient’s clinical stability continued to deteriorate, and he did not survive beyond post-operative day three. mNGS was retrospectively performed on plasma samples collected pre- and post-transfusion from the patient, on aliquots from the blood culture bottles that eventually turned positive, and on residual packed red blood cells (PRBCs) remaining following transfusion. Following receipt of samples from the affected hospital, library preparation, sequencing and preliminary analysis time totaled 72 hours. No remaining material from the platelets or transfused plasma was available for mNGS or other diagnostic testing. Cultured bacterial isolates were unavailable for sequencing.

mNGS revealed no *P. aeruginosa* in the pre-transfusion plasma sample nor in the residual PRBCs, but identified a high abundance of *P. aeruginosa* in post-transfusion plasma (**Figure 2**) that decreased over time in the setting of antibiotic treatment. Serial measurement of procalcitonin demonstrated a normal pre-transfusion level (0.034 ng/mL, reference interval: <0.15 ng/mL) but significantly elevated concentrations following surgery (range: 42.73 to >100 ng/mL [above detection limit]). Phylogenetic analysis indicated strong relatedness to *P. aeruginosa* strain BWHPSA041.

Several environmentally ubiquitous *Pseudomonas* species are common contaminants of mNGS library preparation reagents and thus we used IDSeq[11] to determine the percent of *Pseudomonas* genus reads that mapped specifically to *P. aeruginosa*. In all six post-transfusion plasma samples, as well as the blood culture bottle samples, over 99% of genus *Pseudomonas* reads mapped best to *P. aeruginosa*, while in water controls an average of 7.2% (range 0-12.1%) mapped best to *P. aeruginosa* (**Supplemental Figure 1B**). This result indicated that the *P. aeruginosa* observed in the post-transfusion plasma samples was not the result of contamination from mNGS library preparation reagents.

### Investigation #3: Fatal *Klebsiella* septic platelet transfusion

Two immunocompromised pediatric patients, 2 and 3 years of age, developed septic shock following transfusion of platelets derived from a single donor, as recently reported[6]. Patient C, who had undergone successful autologous hematopoietic stem cell transplantation, developed hypotension, tachycardia and vomiting 15 minutes into the transfusion. Despite initiation of vancomycin and cefepime, fluid resuscitation, vasopressor support and intubation, he died within five hours. Blood cultures ultimately returned positive for *Klebsiella pneumoniae*, as did a culture from residual material in the platelet bag.

Five hours earlier, Patient D, who was receiving empiric cefepime for neutropenic fever, had undergone transfusion with the second platelet unit derived from the same donor. He decompensated into septic shock nine hours following transfusion but ultimately survived following fluid resuscitation and vasopressor support. Blood cultures remained negative in the setting of concurrent antibiotic treatment, initially precluding determination of whether sepsis was due to a Klebsiella-contaminated platelet transfusion or to another etiology.

Culture-independent mNGS of pre- and post-transfusion plasma or serum samples revealed a marked increase in *Klebsiella pneumoniae* following transfusion in both patients, although patient D, who had negative blood cultures, had 12.9-fold fewer *Klebsiella* rpM detected compared to patient C (**Figure 3**). Genome assembly and phylogenetic analysis revealed only one SNP across the 4.2 Mb core genome of *K. pneumoniae* among Patient C’s plasma sample, Patient C’s blood culture isolate, and the residual platelet bag culture isolate. The residual platelet product in the bag given to Patient D was discarded by the hospital nursing staff and unavailable for either culture or mNGS. Across approximately 2700 bases of the *K. pneumoniae* genome detected in Patient D’s plasma with a read depth greater than 10, zero SNPs relative to the other three samples were identified. The sequence type of *K. pneumoniae* in all four samples was determined to be ST491, and the closest ST491 sequence on Genbank differed from these sequences by 212 SNPs.

**Figure 3.**
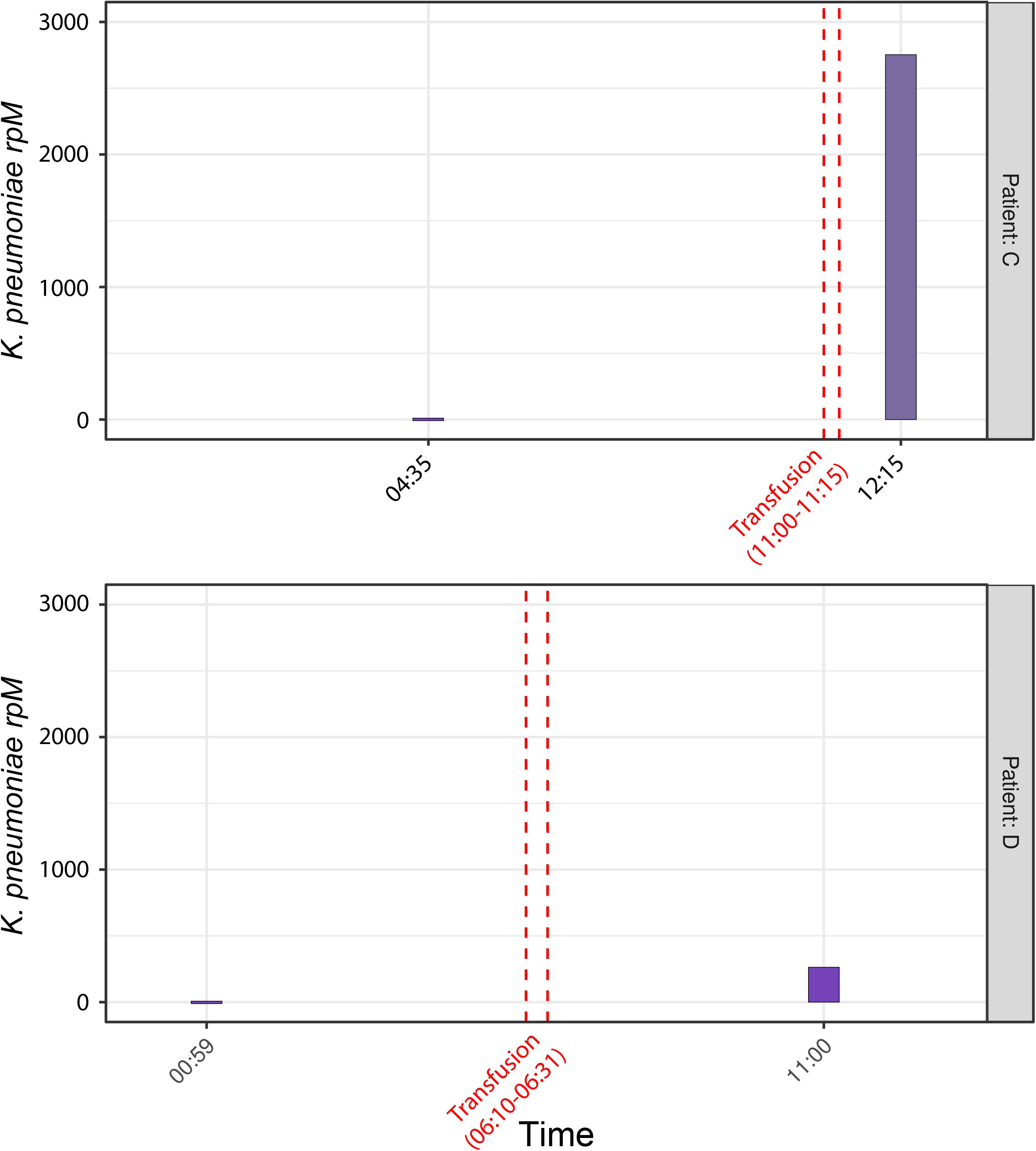
*Klebsiella* septic transfusion investigation. Abundance of *Klebsiella pneumoniae* in plasma from patients C (upper panel) and D (lower panel) during the course of related septic transfusion events, as determined by culture-independent metagenomic sequencing and measured in reads per million (rpM). Patient C, who did not survive the event, had post-transfusion blood cultures return positive for *K. pneumoniae* that was highly related (1 SNP across the 4.2 Mb core genome) to the *K. pneumoniae* isolated from the residual transfused platelet product. Patient D, who was receiving antibiotics with activity against *Klebsiella* prior to transfusion, survived, but had negative post-transfusion blood cultures, precluding definitive confirmation of a related second septic transfusion event in the absence of culture-independent metagenomic sequencing.

Together these data confirmed that the platelet unit represented a single source of infection and that despite Patient D having negative blood cultures, both patients became bacteremic with the same strain of *K. pneumoniae* transfused from the contaminated platelet components. As previously reported, routine culture-based screening of the donor’s platelets at 24 hours performed by the blood supplier remained negative at five days, although an additional platelet unit from the same donor shipped to a different hospital and quarantined before transfusion grew *K. pneumonaie* that was highly related based on whole genome sequencing analysis[6].

## Discussion

Bacterial contamination of platelet products remains an important and underrecognized hospital infection control challenge despite existing screening methods and psoralen-based pathogen-reduction strategies[1–5]. Rapid recognition of potential transfusion-associated sepsis can permit quarantine of untransfused co-components from a potentially contaminated supply chain and assist with root cause analysis. Traditionally, culture-based methods including pulsed-field gel electrophoresis and more recently, whole genome sequencing, have been the central diagnostic tools for septic transfusion investigations. Here, we found that culture-independent mNGS extended the utility of these methods by directly detecting pathogens from clinical samples to assess genetic relatedness, obtain precise strain information, and interrogate levels of pathogen in a patient’s bloodstream throughout the course of a septic transfusion event.

In each case examined, mNGS provided detailed and precise information that clarified the sequence of events resulting in transfusion-related sepsis. In case one, for example, potential explanations considered included: 1) failure of the pathogen inactivation process; 2) contamination of the platelet bag from an environmental source after pathogen reduction; and 3) pre-existing occult bacteremia resulting in retrograde introduction of bacteria into the platelet bag during transfusion. Temporal mNGS assessment of patient plasma demonstrated abundant *Acinetobacter* DNA in post-transfusion samples and in washes of the transfused platelet bag, but not in any pre-transfusion samples, consistent with a septic transfusion event and not preexisting bacteremia.

The findings of high abundance *Acinetobacter sp*. in the transfused unit but not untransfused co-components suggested that contamination occurred following pathogen reduction treatment, potentially from an environmental source during product handling, transport or storage. Transfusion-related sepsis from pathogen-reduced platelet products has been reported but is extremely rare[9,10]. This case suggests that pathogen reduction technology should still be accompanied by rigorous infection control precautions during downstream processing and potentially undergo the same culture-based sterility testing as other platelet products. Discovery that the implicated pathogen in case one represented a novel species of *Acinetobacter* also highlighted the unique ability of sequencing-based diagnostics for unbiased microbe discovery.

In case two, the possibility of an occult but developing bloodstream infection present prior to transfusion was also considered as a potential explanation for the patient’s post-surgical sepsis. As in case one, assessment of plasma samples collected before and after transfusion clarified the sequence of events and demonstrated that *P. aeruginosa* was only detectable post-transfusion. Evidence for a septic transfusion event was further corroborated by temporal measurement of procalcitonin, a host inflammatory biomarker with specificity for bacterial infection[22,23]. Post-transfusion procalcitonin levels above the upper limit of detection in the context of normal pre-surgical levels provided further evidence that a septic transfusion event had occurred. The absence of detectable *P. aeruginosa* in the transfused PRBC segments suggested that platelets or plasma may have been the source of contamination, although neither sample type was available for confirmation.

All three cases notably demonstrated that mNGS afforded high resolution taxonomic identification without a need to isolate a pathogen in culture. This allowed for post-hoc analyses of banked clinical specimens obtained both pre- and post-transfusion that were either unsuitable or unavailable for culture. For instance, even though no cultured isolates were available in case two, direct mNGS of leftover blood products allowed for precise identification of the most closely related *Pseudomonas aeruginosa* strain, which incidentally was recovered in 2013 from a patient’s wound in Massachusetts, USA.

Confirming transmission of a pathogen during septic transfusion events is essential for hospital infection control, but in some cases is not possible because culture fails to identify a microbe. This problem was highlighted by case three, in which blood cultures from patient D remained negative despite the development of post-transfusion septic shock. Patient D was receiving a prophylactic antibiotic with activity against *Klebsiella*, which likely inhibited bacterial growth in culture, precluding definite confirmation of a septic transfusion event related to that experienced by Patient C. Culture-independent mNGS not only confirmed the presence of *K. pneumoniae* in the blood of both patients but also established that it was identical to the isolate derived from transfused platelet product. This unfortunate fatal transfusion case demonstrated the capability of mNGS to provide definitive confirmation and characterization of septic transfusion events in cases where culture fails to yield an isolate.

*Acinetobacter* and *Pseudomonas* are environmentally ubiquitous and common contaminants of mNGS and 16S rRNA gene sequencing library preparation reagents[21,24], and as such can add considerable complexity to investigations in which accurately assessing the abundance of transfused pathogens belonging to these genera is critical. To address this, we employed two complementary approaches that may be broadly useful for future investigations: 1) assessing compositional changes of species within the relevant genus as a proxy for environmental contamination, and 2) focusing analysis exclusively on the exact species implicated in the transfusion event. In case one, for example, these approaches clarified that the novel *Acinetobacter* species was present only in the residual transfused product and in the patient’s plasma following transfusion, but not in the other co-components nor in the patient’s bloodstream prior to the event. In case two, the lack of a *Pseudomonas* cultured isolate limited our ability to perform phylogenetic analyses, but mNGS clearly identified the most closely related species to the causative agent and exhibited a stark compositional difference between the pre- and post-transfusion samples, demonstrating that metagenomic bioinformatics tools like IDSeq can make species-level assignments of individual reads or contigs even in the absence of culture-based whole genome sequencing.

Rapid assessment of septic transfusion events is critical to ensure related contaminated products can be swiftly quarantined, and probable sources of contamination identified. We found that mNGS and pathogen analysis could be reliably performed in under 48 hours, faster than the turnaround time for blood culture at many institutions. Cost, time and infrastructure requirements currently make sequencing impractical at many healthcare institutions, however new platforms such as the Illumina iSeq and Oxford Nanopore Minion will undoubtedly increase the broad applicability of this technology for rapid hospital epidemiologic investigations.

As genomic approaches become more widely used for investigating transfusion-related infections, rapid exchange of pathogen genomic information via open access databases could accelerate identification of related cases, enhancing infection control efforts of emerging outbreaks. Indeed, the findings described here have contributed to a multicenter United States Centers for Disease Control and Prevention investigation which has identified the novel *Acinetobacter sp*. from Patient A in related cases from Utah and Connecticut[25].

In summary, transfusion-related sepsis continues to cause excess mortality and morbidity despite the introduction of pathogen-reduction technologies. We found that culture-independent mNGS complemented current best available methods for investigation of transfusion-related sepsis by extending traditional whole genome sequencing-based phylogenetics of cultured isolates and by permitting longitudinal assessment of pathogen abundance pre- and post-transfusion from direct clinical specimens. While additional studies are needed to validate these methods, implementation of mNGS for both investigation and prevention of transfusion-related infections may enhance existing practices.

## Data Availability

Raw sequencing data is available via NCBI BioProject Accession ID: PRJNA544865.

## Acknowledgements

We thank Norma Neff for her guidance on metagenomic sequencing methodology and PK for his support and input on the study. We thank Amy Kistler and Renuka Kumar for their efforts with data analysis and sample processing. We thank Michael Busch for sharing his expertise in evaluating transfusion-transmitted infections.

## Authorship Contributions

C.L., E.C., J.K. and S.M. wrote the paper

J.K. and L.L. performed phylogenetic analyses

C.L., E.C., J.K., L.L. and PC performed additional data analyses

A.N., A.N., D.Y., M.M., M.Z., N.T. and S.B. directed clinical aspects of the investigations

C.T. and J.D. directed molecular aspects of the investigations

A.L., B.P., C.L., E.C., P.H., and J.Q. performed sample extractions and metagenomic library preparation

M.P. served as clinical research coordinator

M.T. and R.S. performed sequencing and quality control

## Supplemental Methods

### Sample Processing and Nucleic Acid Extraction

Samples were inactivated by adding DNA/RNA Shield (Zymo) and bashed with ceramic beads on a TissueLyser (Qiagen) for either two minutes at 15s^-1^ or one minute at 30s^-1^. For case one, metagenomic samples were extracted with the Quick-DNA/RNA Miniprep Kit (Zymo) and cultured isolate samples were extracted with the Quick DNA Fungal/Bacterial kit (Zymo). Case two samples were extracted with the Quick-DNA/RNA Microprep Plus kit (Zymo). For case three, metagenomic samples were extracted with the Duet DNA/RNA kit (Zymo) and cultured isolate samples were extracted with the AllPrep DNA/RNA kit (Qiagen) on a Qiacube robot.

### Sterility Testing

The United States Food and Drug Administration (FDA) does not require primary culture testing for pathogen-reduced platelet products, as such no sterility testing was done for the platelet products from case 1. For the remaining cases, apheresis platelet products were subjected to sterility testing by the blood supplier using a standard FDA-approved automated culture methods and protocol. This involved culture performed by the blood supplier 24 hours after collection of the donor’s platelets. Eight mL of platelet product was inoculated into an aerobic BacT bottle and incubated at 37 degrees Celsius for 5 days. In case 3, The BacT bottle remained negative at the end of 5 days of incubation. False negativity is a known limitation of culture-based platelet sterility testing and may reflect low levels of inoculum at the time of platelet collection that are not picked up by sampling a small volume of product. Transfusion services are not currently required to perform additional cultures or point of release tests for bacterial contamination prior to issuing platelets to patients. Plasma products are not subjected to sterility testing by the blood supplier or by the transfusion service.

**Supplemental Table 1:** Microbial Culture Data.

| **Case** | **Pathogen** | **Culture type** | **Time to positivity** | **Time to AST reporting** |
| --- | --- | --- | --- | --- |
| 1 | *Acinetobacter* | Product initial wash | 14h 41m | 66h 39m |
| 1 | *Acinetobacter* | Product repeat wash | 6h 58m | 59h 46m |
| 1 | *Acinetobacter* | Central blood | 15h 54m | 66h 4m |
| 1 | *Acinetobacter* | Peripheral blood | 24h 34m | 135h 41m |
| 2 | *Pseudomonas* | Central blood | 18h 19m | 59h 22m |
| 2 | *Pseudomonas* | Peripheral blood | 27h 14m |  |
| 3 | *Klebsiella* | Central blood red port | 7h 6m |  |
| 3 | *Klebsiella* | Central blood white port | 5h 6m | 42h 16m |
| 3 | *Klebsiella* | Peripheral blood | 95h 7m |  |
| 3 | *Klebsiella* | Platelet product | 1h 6m | 62h 21m |

Legend: AST = antimicrobial susceptibility testing.

**Supplemental Table 2:** Sequencing reads collected per sample.

| Case | Sample Name | Day | Time | Total Reads | Pathogen Reads | Pathogen |
| --- | --- | --- | --- | --- | --- | --- |
| 1 | H2O-1a | n/a | n/a | 12292 | 0 | *ACB complex* |
| 1 | H2O-1b | n/a | n/a | 4196 | 0 | *ACB complex* |
| 1 | Co-component-1 | n/a | n/a | 33437780 | 87 | *ACB complex* |
| 1 | Co-component-2 | n/a | n/a | 105027102 | 190 | *ACB complex* |
| 1 | Patient A | Day (-1) | 5:01 | 111863490 | 58 | *ACB complex* |
| 1 | Patient A | Day 0 | 3:17 | 91789296 | 23 | *ACB complex* |
| 1 | Patient A | Day 1 | 4:52 | 86230676 | 91 | *ACB complex* |
| 1 | Patient A | Day 1 | 14:41a | 89397082 | 19092 | *ACB complex* |
| 1 | Patient A | Day 1 | 14:41b | 150000000 | 30666 | *ACB complex* |
| 1 | Patient A | Day 1 | 15:23 | 148209522 | 21096 | *ACB complex* |
| 1 | H2O-2a | n/a | n/a | 5532686 | 1290 | *ACB complex* |
| 1 | Transfused platelets | n/a | n/a | 150000000 | 34367349 | *ACB complex* |
| 1 | Isolate-a (Platelet bag) | n/a | n/a | 10273280 | - | *ACB complex* |
| 1 | isolate-b (Platelet bag) | n/a | n/a | 7746330 | - | *ACB complex* |
| 1 | isolate-c (Patient) | n/a | n/a | 6458218 | - | *ACB complex* |
| 2 | H2O-1a | n/a | n/a | 56452 | 3 | *P. aeruginosa* |
| 2 | H2O-1b | n/a | n/a | 1427720 | 155 | *P. aeruginosa* |
| 2 | H2O-1c | n/a | n/a | 283232 | 32 | *P. aeruginosa* |
| 2 | Blood culture bottle-1 | Day 2 | 16:06 | 150000000 | 1223 | *P. aeruginosa* |
| 2 | Blood culture bottle-2 | Day 2 | 16:06 | 150000000 | 1533 | *P. aeruginosa* |
| 2 | H2O-2a | n/a | n/a | 9523090 | 399 | *P. aeruginosa* |
| 2 | Patient B | Day 1 | 3:55 | 150000000 | 5 | *P. aeruginosa* |
| 2 | Patient B | Day 1 | 21:09 | 150000000 | 81995 | *P. aeruginosa* |
| 2 | Patient B | Day 2 | 3:49 | 118341948 | 4248 | *P. aeruginosa* |
| 2 | Patient B | Day 2 | 11:36 | 99059392 | 875 | *P. aeruginosa* |
| 2 | Patient B | Day 2 | 21:44 | 62467588 | 269 | *P. aeruginosa* |
| 2 | Patient B | Day 2 | 23:19 | 83908304 | 178 | *P. aeruginosa* |
| 2 | Patient B | Day 3 | 1:38 | 110713036 | 87 | *P. aeruginosa* |
| 2 | PRBC-1 | n/a | n/a | 150000000 | 24 | *P. aeruginosa* |
| 2 | PRBC-2 | n/a | n/a | 150000000 | 234 | *P. aeruginosa* |
| 3 | Patient C | Day 1 | 4:35 | 125893204 | 31 | *K. pneumoniae* |
| 3 | Patient C | Day 1 | 12:15 | 45615102 | 1255715 | *K. pneumoniae* |
| 3 | Patient D | Day 1 | 0:59 | 10769438 | 42 | *K. pneumoniae* |
| 3 | Patient D | Day 1 | 11:00 | 42995136 | 113123 | *K. pneumoniae* |
| 3 | H2O-1a | n/a | n/a | 2734220 | 217 | *K. pneumoniae* |
| 3 | Isolate-a (Patient C) | n/a | n/a | 6601096 | - | *K. pneumoniae* |
| 3 | Isolate-b (Platelet bag) | n/a | n/a | 5800012 | - | *K. pneumoniae* |

**Supplemental Figure 1.**
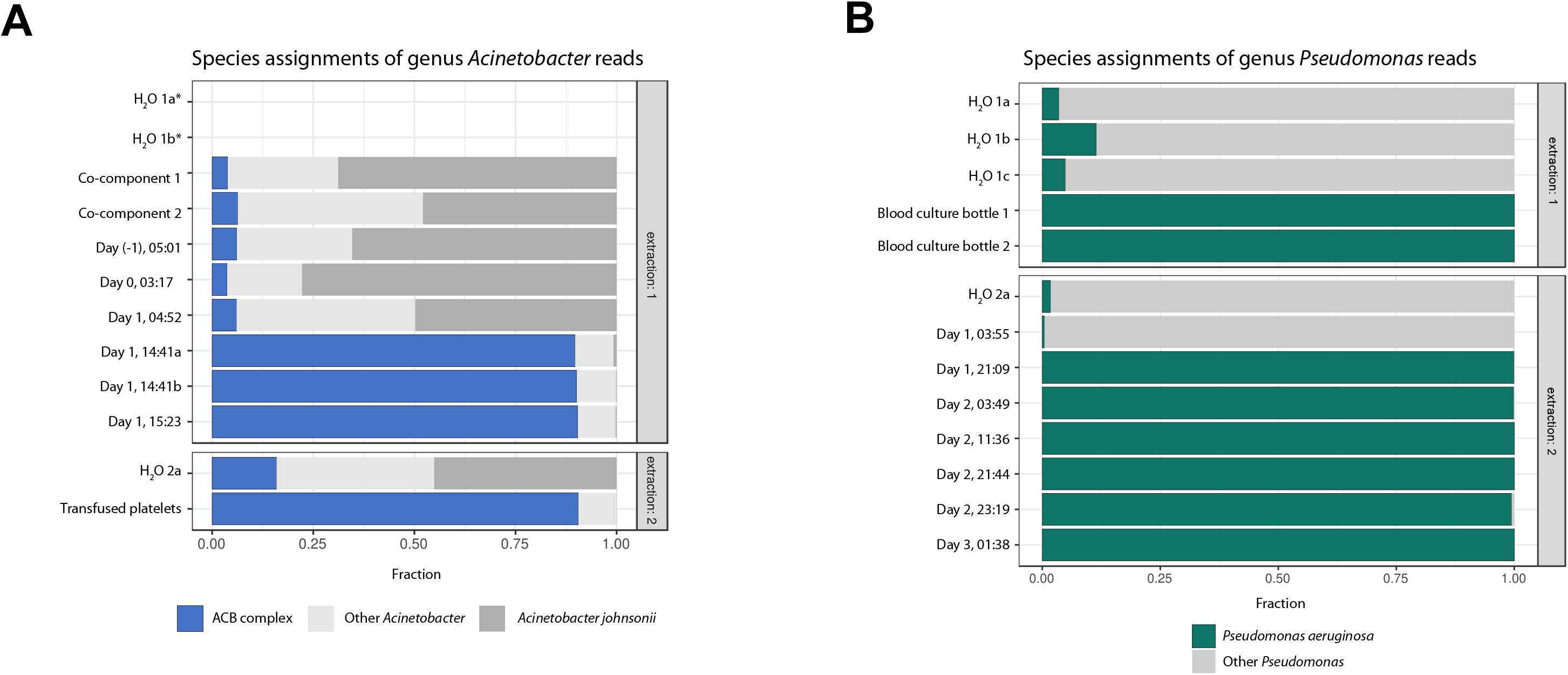
Fraction of species-specific sequencing reads aligning to transfusion-transmitted pathogens. **A)** Fraction of species-specific *Acinetobacter* read assignments for samples evaluated by culture-independent metagenomic sequencing in case 1, including water controls. The cocomponents and pre-transfusion samples (upper panel) had only a small fraction of genus *Acinetobacter* reads that best aligned to *Acinetobacter calcoaceticus/baumannii* (ACB) complex while the transfused product (lower panel) and patient’s post-transfusion plasma (upper panel) was comprised mostly of reads aligning to the ACB complex. Asterisks indicate samples with no reads to genus *Acinetobacter*. **B)** Fraction of species-specific *Pseudomonas* sequencing read assignments for plasma samples evaluated by culture-independent metagenomic sequencing for case 2, including water controls. Samples from the blood culture bottles which eventually turned positive (upper panel) and from the patient’s plasma post-transfusion (lower panel) were comprised almost entirely of reads best aligning to *Pseudomonas aeruginosa*, while the patient’s plasma before transfusion and water control samples had mostly reads aligning to other *Pseudomonas* species.

## References

1. Eder AF, Dy BA, DeMerse B, et al. Apheresis technology correlates with bacterial contamination of platelets and reported septic transfusion reactions: APHERESIS TECHNOLOGY AND BACTERIAL CONTAMINATION OF PLTs. Transfusion (Paris) 2017; 57:2969–2976.

2. Kaufman RM, Assmann SF, Triulzi DJ, et al. Transfusion-related adverse events in the Platelet Dose study. Transfusion (Paris) 2015; 55:144–153.

3. Busch MP, Bloch EM, Kleinman SH. Prevention of transfusion transmitted infections. Blood 2019;

4. Tormey CA, Sweeney JD, Champion MH, Pisciotto PT, Snyder EL, Wu Y. Analysis of transfusion reactions associated with prestorage-pooled platelet components. Transfusion (Paris) 2009; 49:1242–1247.

5. Kuehnert MJ, Roth VR, Haley NR, et al. Transfusion-transmitted bacterial infection in the United States, 1998 through 2000. Transfusion (Paris) 2001; 41:1493–1499.

6. Horth RZ, Jones JM, Kim JJ, et al. Fatal Sepsis Associated with Bacterial Contamination of Platelets — Utah and California, August 2017. MMWR Morb Mortal Wkly Rep 2018; 67:718–722.

7. Fuller AK, Uglik KM, Savage WJ, Ness PM, King KE. Bacterial culture reduces but does not eliminate the risk of septic transfusion reactions to single-donor platelets. Transfusion (Paris) 2009; 49:2588–2593.

8. Lin L, Dikeman R, Molini B, et al. Photochemical treatment of platelet concentrates with amotosalen and long-wavelength ultraviolet light inactivates a broad spectrum of pathogenic bacteria. Transfusion (Paris) 2004; 44:1496–1504.

9. Schmidt M, Hourfar MK, Sireis W, et al. Evaluation of the effectiveness of a pathogen inactivation technology against clinically relevant transfusion-transmitted bacterial strains: EFFICIENCY OF PATHOGEN INACTIVATION. Transfusion (Paris) 2015; 55:2104–2112.

10. Benjamin RJ, Braschler T, Weingand T, Corash LM. Hemovigilance monitoring of platelet septic reactions with effective bacterial protection systems: HEMOVIGILANCE FOR TRANSFUSION SEPSIS. Transfusion (Paris) 2017; 57:2946–2957.

11. Ramesh A, Nakielny S, Hsu J, et al. Etiology of fever in Ugandan children: identification of microbial pathogens using metagenomic next-generation sequencing and IDseq, a platform for unbiased metagenomic analysis. 2018; Available at: http://biorxiv.org/lookup/doi/10.1101/385005. Accessed 3 November 2018.

12. TrimGalore. Available at: https://github.com/FelixKrueger/TrimGalore.

13. Wick RR, Judd LM, Gorrie CL, Holt KE. Unicycler: Resolving bacterial genome assemblies from short and long sequencing reads. PLOS Comput Biol 2017; 13:e1005595.

14. Camacho C, Coulouris G, Avagyan V, et al. BLAST+: architecture and applications. BMC Bioinformatics 2009; 10:421.

15. Ondov BD, Treangen TJ, Melsted P, et al. Mash: fast genome and metagenome distance estimation using MinHash. Genome Biol 2016; 17. Available at: http://genomebiology.biomedcentral.com/articles/10.1186/s13059-016-0997-x. Accessed 21 March 2019.

16. Wattam AR, Abraham D, Dalay O, et al. PATRIC, the bacterial bioinformatics database and analysis resource. Nucleic Acids Res 2014; 42:D581–D591.

17. Inouye M, Dashnow H, Raven L-A, et al. SRST2: Rapid genomic surveillance for public health and hospital microbiology labs. Genome Med 2014; 6:90.

18. Snippy. Available at: https://github.com/tseemann/snippy/releases.

19. bcftools v1.9. Available at: https://github.com/samtools/bcftools.

20. Stamatakis A. RAxML version 8: a tool for phylogenetic analysis and post-analysis of large phylogenies. Bioinforma Oxf Engl 2014; 30:1312–1313.

21. Wilson MR, O’Donovan BD, Gelfand JM, et al. Chronic Meningitis Investigated via Metagenomic Next-Generation Sequencing. JAMA Neurol 2018; 75:947–955.

22. Vijayan AL, Vanimaya null, Ravindran S, et al. Procalcitonin: a promising diagnostic marker for sepsis and antibiotic therapy. J Intensive Care 2017; 5:51.

23. Liu D, Su L, Han G, Yan P, Xie L. Prognostic Value of Procalcitonin in Adult Patients with Sepsis: A Systematic Review and Meta-Analysis. PLOS ONE 2015; 10:e0129450.

24. Salter SJ, Cox MJ, Turek EM, et al. Reagent and laboratory contamination can critically impact sequence-based microbiome analyses. BMC Biol 2014; 12. Available at: 3https://bmcbiol.biomedcentral.com/articles/10.1186/s12915-014-0087-z. Accessed 10 May 2019.

25. Jones SA, Jones, JM, Leung V, Nakashima AK, Oakeson KF, Simth AR, Hunter R, Kim JJ, Cumming M, McHale E, Young PP, Fridey JL, Kelley WE, Stramer SL, Wagner SJ, West FB, Heron R, Snyder E, Hendrickson J, Peaper DR, Gundlapalli AV, Langelier C, Miller S, Nambiar A, Moayeri M, Kamm J, Moulton-Meissner, H, Annambhotla P, Gable P, McAllister GA, Breaker E, Sula E, Halpin AL, Basavaraju SV. Septic Transfusion Reactions Attributed to Bacterial Contamination of Platelets Associated with a Potential Common Source — Multiple States, 2018. MMWR Morb Mortal Wkly Rep 2019;

